# Extracellular vesicles isolated from frozen and fresh human melanoma tissue are similar in purity and protein composition

**DOI:** 10.1101/2024.04.03.587936

**Authors:** Daniele D’Arrigo, Cecilia Lässer, Ornella Urzì, Kyong-Su Park, Roger Olofsson Bagge, Jan Lötvall, Rossella Crescitelli

## Abstract

Extracellular vesicles (EVs) isolated from tumor tissues represent a precious source of EVs because they are enriched in disease-associated biomarkers and are highly concentrated. However, a fundamental question is whether freezing of tissues influences the EVs’ integrity and function and whether non-EVs are co-isolated with the EVs. In this work, we isolated EVs from metastatic melanoma tissue both immediately after tissue resection and after being frozen for at least two weeks. Specifically, the samples were divided into two parts: one was immediately processed for EV isolation (hereafter called fresh), and the other was frozen on dry ice and stored for two weeks before being processed for EV isolation (hereafter called frozen). Large and small EVs were isolated through ultracentrifugation, pooled, and further purified on an iodixanol density cushion. The EVs were analyzed by transmission electron microscopy, nanoparticle tracking analysis, and protein quantification as well as by quantitative mass spectrometry. The results did not show any significant difference between EVs isolated from fresh and frozen tissue. Importantly, there was no enrichment of either intracellular proteins or mitochondrial proteins in EVs isolated from frozen tissues vs. fresh tissues. Moreover, there were no significant differences in the quantity of proteins like MT-CO2, Cox6, SLC24A22, HLA-DR, and Erlin2, which were previously identified as potential markers of melanoma, ovarian cancer, and breast cancer. Overall, this study supports the use of frozen tissues as a source of EVs for research purposes because frozen tissue-derived EVs do not differ in purity or protein composition compared to their fresh counterparts.

## 1. Introduction

A complex network of interactions between cells is required for the maintenance of tissue homeostasis ^1^. In these processes, extracellular vesicles (EVs) represent one key player in cell-to-cell communication because they convey complicated signaling between cells, including membrane proteins and intravesicular RNA or proteins ^2–4^. EVs are also implicated in the progression of multiple types of tumors ^5^ and are emerging as possible disease biomarkers because they can reflect the pathophysiological state of the EV-producing cell ^6^.

The first and more widespread approach for exploiting the diagnostic clues associated with EVs is based on the analysis of EVs in biofluids, including blood, urine, and saliva ^7, 8^. This approach has some advantages like a relatively non-invasive collection procedure ^8^; however it comes with some limitations. Despite the efforts of the scientific community, EVs isolated from biofluids have shown low diagnostic efficacy, low robustness, and lack of specificity for several diseases ^8–11^. In addition, EVs are extremely diluted in biofluids, making their detection challenging. In contrast, EVs isolated from solid tissues may better resemble the diseased microenvironment ^12, 13^. To date, this approach has been considerably less exploited compared to the analysis of liquid biopsies, and only recently have protocols for EV isolation from solid tissues been developed ^12, 14–17^. Both fresh and frozen tissues have been used as the source of EVs, but cryopreserved samples undergo greater manipulation that can lead to a potentially greater risk of cell membrane damage and to the release of intracellular vesicles ^18^. Importantly, a large number of samples from existing biobanks could potentially be used as source of EVs, thus greatly increasing the possibility for developing EV-based diagnostic tools.

The detailed freezing procedures are not reported in many studies focusing on tissue-derived EVs, but the effects of the freezing procedure on EV preparations need to be properly investigated because the freezing process can lead to cell disruption ^18^ and consequently to contamination of EV samples by intracellular vesicles ^19^. Different protocols for freezing solid tissues have been developed, each with its advantages and disadvantages. Snap freezing in isopentane or in liquid nitrogen and direct freezing on dry ice or in a –80°C freezer are the most common ways to preserve tissue samples ^20–22^. The last procedure has a high risk of causing cryo-artifacts in the tissue, but at the same time it represents an easier, faster, and less biohazardous approach compared to other approaches ^22^. In addition, –80°C is adequate for the storage of tissues even for extended time periods ^23, 24^ and it has primarily been employed in works involving EVs from frozen tissues ^16, 17, 25–30^, thus representing a feasible approach for most laboratories.

Despite the number of studies focused on EVs isolated from frozen tissues, they have only investigated the presence of a few intra-cellular markers to ensure the purity of tissue-derived EVs ^13, 16, 17, 25–33^. In this study we analyzed the purity, the proteomic content, and the maintenance of the diagnostic potential of EVs from frozen tissue in comparison to fresh tissue-derived EVs.

## 2. Materials and methods

### 2.1 Patient information

Melanoma tissues were collected from five patients with a confirmed diagnosis of stage III or IV melanoma during surgical resection at the Department of Surgery at Sahlgrenska University Hospital between December 2021 and February 2022. The biological material collected for this study included two cutaneous metastases and three lymph node metastases with a weight ranging from 0.2 g to 3.9 g. Patient demographics are described in Table 1. Ethics permission was granted by the Regional Ethical Review Board at the University of Gothenburg, Sweden (Dnr #096-12 and 995-16). The patients provided written informed consent.

**TABLE 1.** Patient demographic information

| Patient | Type of metastasis | Sex | Age | TNM-stage | Sample used for |
| --- | --- | --- | --- | --- | --- |
| 1 | Cutaneous metastasis (in-transit) | Male | 59 | Stage IIIC | Cell analysis: single cell isolation, cytospin, EV analysis: TEM, NTA, LC-MS/MS |
| 2 | Lymph node | Male | 48 | Stage IV | Cell analysis: single cell isolation, cytospin, EV analysis: NTA |
| 3 | Lymph node | Male | 56 | Stage IV | Cell analysis: single cell isolation, cytospin, EV analysis: WB, ELISA, NTA, LC-MS/MS |
| 4 | Skin metastasis | Male | 56 | Stage IV | Cell analysis: single cell isolation, cytospin, EV analysis: TEM, NTA, LC-MS/MS |
| 5 | Lymph node | Male | 81 | Stage IIID | Cell analysis: single cell isolation, cytospin, EV analysis: WB, ELISA, NTA, LC-MS/MS |

TEM= Transmission Electron Microscopy
NTA= Nanoparticle Tracking Analysis
LC-MS/MS= Liquid Chromatography Tandem Mass Spectrometry

### 2.2 Tissue handling and freezing

Immediately after the tumor resection, the samples were weighed and divided in two equal portions. Both portions were placed in RPMI-1640 (Sigma-Aldrich, St Louis, MO) and cut into pieces of approximately 2 mm × 2 mm × 2 mm with a scalpel. One portion was immediately processed (fresh samples), while the other portion was frozen on dry ice (frozen samples). The latter was put in 1.5 mL conical cryovials to minimize the formation of air bubbles as much as possible. The cryovials were then placed in a metallic rack (Corning, Corning, NY) that had been cooled in an ice pan filled with dry ice and incubated until completely frozen. The frozen tissues were kept for 2 weeks at –80°C until further processing. The procedure is shown in Fig. 1.1.

**FIGURE 1.**
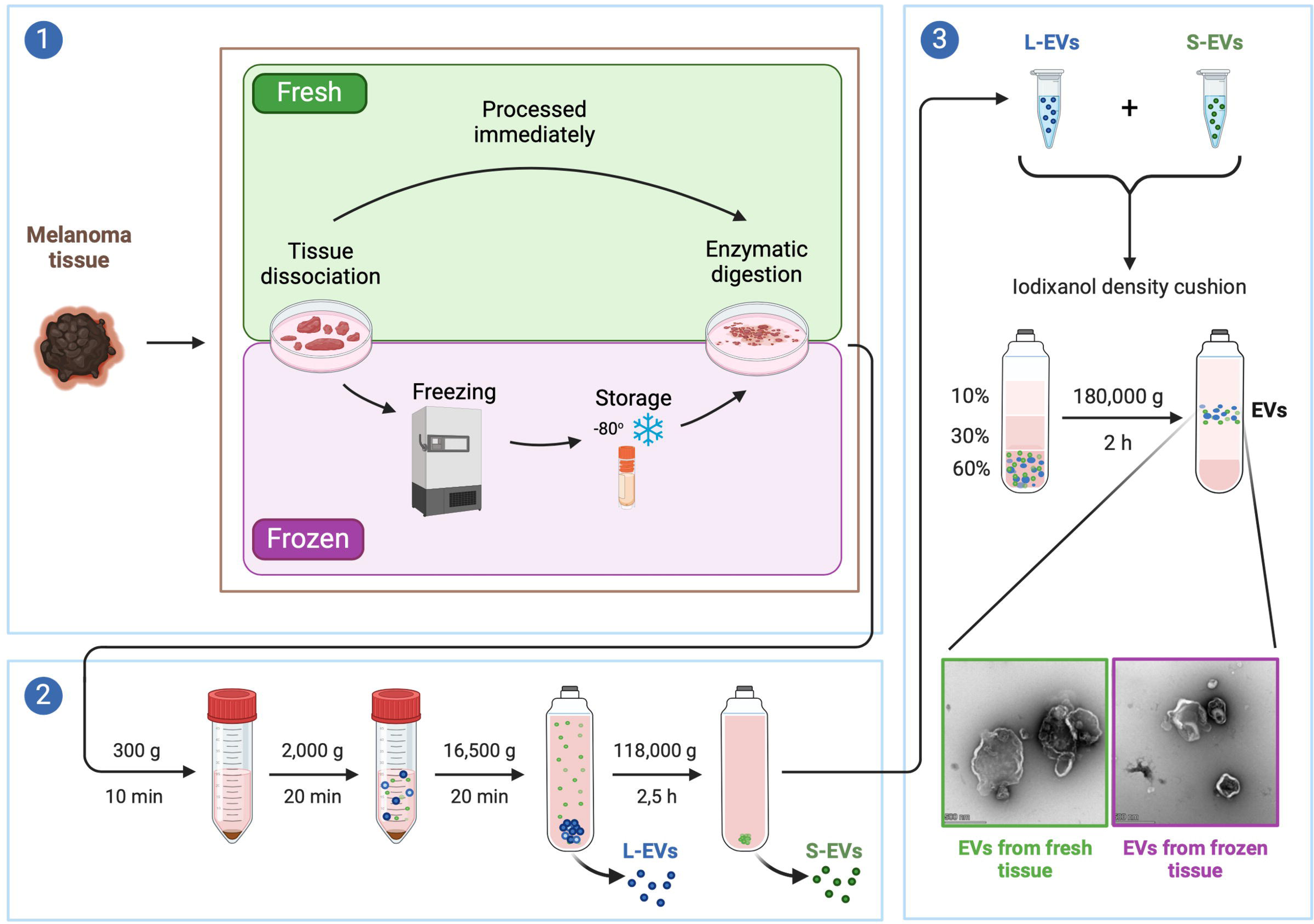
Graphical representation of the experimental design. The melanoma tissues were divided in two parts after collection. One part was immediately processed to isolate EVs. The second part was dissected into small tissue pieces and frozen in dry ice and stored at –80°C for two weeks before being processed for EV isolation (1) through a combination of differential centrifugations (2) and density cushion (3).

### 2.3 EV isolation from human melanoma metastatic tissue

EVs were immediately isolated from the fresh tissue, while the frozen sample was frozen for two weeks and then thawed on ice before being processed. The same protocol, described by Crescitelli *et al* ^15^, was used for isolating EVs from both the fresh and frozen tissues. In brief, just after the resection the tissue was dissociated into small pieces (∼2 mm × 2 mm × 2 mm) and then digested for 30 minutes at 37°C with Collagenase (2 mg/mL, Roche, Basel, Switzerland) and DNase I (400 U/mL, Roche). The digested tissue pieces were then filtered through a 70 µm filter. The flow-through was used for the EV isolation (see paragraph 2.4), while the parts remining in the filter (tissue pieces collected after enzymatic treatment and filtration) were further processed as described in paragraph 2.5. The procedure is schematized in Fig. 1. Potential contaminants by cells and cell debris were removed by two centrifugation steps (300 × *g* for 10 min and 2,000 × *g* for 20 min). The supernatant was then transferred to a polypropylene Quick Seal centrifuge tube (Beckman Coulter, Brea, US) and centrifuged at 16,500 × *g* (14,500 rpm) at 4°C for 20 min using a Type Ti 45 fixed-angle rotor (Beckman Coulter, k-factor 1.275) to isolate large EVs. The pellet was then resuspended in PBS and stored at –80°C until further processing, while the supernatant was further centrifuged at 118.000 × *g* (38,800 rpm) (Type Ti 45 fixed-angle rotor, Beckman Coulter, k-factor 178) for 2.5 hours at 4°C to isolate small EVs that were resuspended in PBS and stored at –80°C.

### 2.4 Iodixanol density cushion

EVs isolated by ultracentrifugation were further purified by iodixanol density cushion centrifugation as described by Crescitelli *et al* ^15^. Both large and small EVs were thawed on ice and combined together. The EVs were then mixed with 3 mL of 60% Optiprep (Sigma Aldrich) at the bottom of an Ultra-Clear centrifuge tube (Beckman Coulter), and 4 ml of 30% Optiprep and 4 ml 10% Optiprep were laid on top. Samples were ultracentrifuged at 180,000 × *g* (38,000 rpm) in a SW 41 Ti Swinging-Bucket Rotor (Beckman Coulter, k-factor 145) for 2 hours at 4°C. The samples were collected from the interface of the 10% and 30% iodixanol layers and stored at –80°C.

### 2.5 Tissue dissociation and single cell isolation

The tissue pieces remaining after enzymatic treatment and filtration of the fresh sample were processed using a human tumor dissociation kit (Miltenyi Biotec, Bergisch Gladbach, Germany) according to the manufacturer’s instructions to obtain a single-cell suspension. Briefly, the samples were transferred to a plastic tube containing a knife and placed on the gentleMACS Dissociator (Miltenyi Biotec) to be further dissociated and then incubated with enzymes for 30 min at 37°C under continuous rotation. These steps were repeated three times, and then the dissociated samples were filtered through a 70 µm filter. After centrifugation at 300 × *g* for 7 minutes, the cell pellet was resuspended in RPMI-1640 (Sigma-Aldrich). The cell concentration was assessed using the trypan blue exclusion method.

### 2.6 Flow cytometry analysis of cells isolated from fresh metastatic melanoma tissues

The cellular composition of the solid tumor tissues was determined by flow cytometry analysis. The single cells were incubated with human IgG (Sigma-Aldrich) at 4°C for 15 min to block nonspecific binding sites. The cells were then stained with a viability stain kit (LIVE/DEAD Fixable Aqua Dead Cell Stain Kit, Invitrogen, Life Technologies Corp, Eugene, OR) and antibodies directed against cell-specific surface antigens, namely anti-CD3-APC (1:5 dilution, clone SK7) and anti-CD45-PerCP (1:10 dilution, clone 2D1) (both from BD Biosciences, San Jose, CA). After incubation for 30 min at 4°C in the dark, the cells were washed in wash buffer (1% FBS in PBS) followed by a fixation and permeabilization step with the eBioscience Foxp3/Transcription Factor Staining Buffer Set (Invitrogen) for 1 hour at 4°C in the dark. Antibodies against SOX-10-PE (1:250 dilution, clone SP267) or isotypic control-PE (1:250 dilution, clone EPR25A) (both from Abcam, Cambridge, UK) were added and incubated for 1 hour at 4°C in the dark. After washing, the cells were analyzed on a BD FACSVerse Flow Cytometer equipped with BD FACSSuite software (BD Biosciences), and the data were analyzed with FlowJo software (Tree Star Inc., Ashland, OR). The gating of the signal from each antibody was set by using control samples obtained with the “fluorescence minus one” approach in which the control sample of each antibody contained all markers except the one of interest. Only live cells were included in the gating process. In addition, 50,000 cells were deposited in a single layer on a glass microscope slide using a cytocentrifuge, and they were stained with Hemacolor Rapid stain (Merck Millipore, Darmstadt, Germany) following the manufacturer’s protocol.

### 2.7 Particle measurement

The number of isolated particles was determined using a ZetaView PMX110 instrument (Particle Metrix, Meerbusch, Germany). The measurements were carried out in 11 positions with the video quality set to medium and the sensitivity of the camera set to 80. The temperature of the sample chamber was automatically assessed and integrated into the calculation by the software. Data were analyzed using ZetaView analysis software (version 8.2.30.1) with a minimum and a maximum size of 5 and 5,000, respectively, and a minimum brightness of 20.

### 2.8 Protein measurement

The EVs resuspended in PBS were used to measure the EV protein concentration using a Qubit device (Thermo Fisher Scientific, Waltham, MA) following the manufacturer’s recommendations.

### 2.9 Western blot analysis

For the western blot analysis, proteins were extracted using RIPA buffer (Thermo Fisher Scientific), and 10 µg were then loaded and separated on precast 4–20% polyacrylamide Mini-PROTEAN TGX gels (Bio-Rad Laboratories, Hercules, CA). The proteins were then transferred to PVDF membranes (Bio-Rad Laboratories) that were incubated in EveryBlot Blocking Buffer (Bio-Rad Laboratories) for 5 min at RT to block nonspecific binding. The following primary antibodies diluted in EveryBlot Blocking Buffer (Bio-Rad Laboratories) were then added to the membrane and incubated overnight at 4°C: anti-Calnexin (1:1,000 dilution, clone C5C9, Cell Signaling Technology, Danvers, MA), anti-Mitofilin (1:500 dilution, clone AB-2547893, Invitrogen), anti-ADAM10 (1:500 dilution, clone 163003, R&D System, McKinley Place, MN, in not reducing conditions), anti-CD63 (1:1,000 dilution, clone H5C6, BD Biosciences, not in reducing conditions), anti-Flotillin-1 (1:1,000 dilution, clone EPR6041, Abcam), anti-CD9 (1:1.000 dilution, clone MM2/57, Merck-Millipore, not in reducing conditions), anti-CD81 (1:1,000 dilution, clone M38, Abcam, not in reducing conditions), anti-MTCO2 (1:1,000 dilution, polyclonal, Abcam), and anti-Cox6 (1:500 dilution, clone 3G5F7G3, Abcam). After three washes, the secondary antibodies, diluted in EveryBlot Blocking Buffer (Bio-Rad Laboratories), were added to the membrane and incubated for 1 h at room temperature. The secondary antibodies were anti-rabbit IgG (horseradish peroxidase conjugated, 1:5,000 dilution, Harlan Sera-Lab, Loughborough, UK) and anti-mouse IgG (horseradish peroxidase conjugated, 1:5,000 dilution, Harlan Sera-Lab). The membranes were finally imaged and analyzed with the SuperSignal West Femto maximum sensitivity substrate (Thermo Fisher Scientific) and a ChemiDoc Imaging System (Bio-Rad Laboratories).

### 2.10 ELISA

For the direct ELISA assays, each EV formulation (0.2 μg per well) was coated on a black 96-well plate overnight at 4°C. The plate was blocked with 1% BSA in PBS for 1 h and then incubated with anti-MTCO2 (1:200 dilution, clone 12C4F12, Thermo Fisher Scientific) or anti-Cox6 (1:200 dilution, clone 3G5F7G3, Abcam) diluted in 1% BSA in PBS for 2 h. After washing, the HRP-conjugated anti-mouse antibody (horseradish peroxidase conjugated, 1:5,000 dilution, Harlan Sera-Lab) was diluted in 1% BSA in PBS and incubated for 1 h at room temperature. Luminescent signal was detected with the BM Chemiluminescence ELISA Substrate (BD Biosciences, San Jose, CA).

### 2.11 Transmission electron microscopy (TEM)

EV morphology was investigated by negative staining as previously described ^14, 15^. Briefly, 5 µg of the sample was placed onto a glow discharged 200-mesh formvar/carbon copper grid (Electron Microscopy Sciences, Hatfield Township, PA) for 5 min. After two washes with water, the EV samples were fixed in 2.5% glutaraldehyde and washed again before being stained. To stain the specimens, 2% uranyl acetate was incubated for 1.5 min and the negative-stained EVs were examined on a Talos L120C electron microscope (Thermo Fisher Scientific) at 120 kV with a CCD camera.

### 2.12 Sample preparation and digestion for proteomic analysis

The samples and the reference pool (a representative reference containing aliquots from all the samples) were processed using a modified filter-aided sample preparation (FASP) method ^34^. In short, samples (40 µg) were reduced with 100 mM dithiothreitol at 60°C for 30 min, transferred to Microcon-30kDa Centrifugal Filter Units (Merck), and washed several times with 8 M urea and once with digestion buffer (50 mM TEAB, 0.5% sodium deoxycholate) prior to alkylation with 10 mM methyl methanethiosulfonate in digestion buffer for 30 min at room temperature. Samples were digested with trypsin (Pierce MS-grade trypsin, Thermo Fisher Scientific, 1:100 ratio) at 37°C overnight, and an additional portion of trypsin was added and incubated for another 2 h. Peptides were collected by centrifugation and labelled using tandem mass tag (TMT) 11-plex isobaric mass tagging reagents (Thermo Fisher Scientific) according to the manufacturer’s instructions. The samples were combined into one TMT set, and sodium deoxycholate was removed by acidification with 10% TFA. The TMT set was further purified using High Protein and Peptide Recovery Detergent Removal spin columns and Pierce peptide desalting spin columns (both from Thermo Fischer Scientific) according to the manufacturer’s instructions prior to basic reversed-phase chromatography (bRP-LC) fractionation. Peptide separation was performed using a Dionex Ultimate 3000 UPLC system (Thermo Fischer Scientific) and a reversed-phase XBridge BEH C18 column (3.5 μm, 3.0 × 150 mm, Waters Corporation, Milford, Massachusetts, MA) with a gradient from 3% to 100% acetonitrile in 10 mM ammonium formate at pH 10.00 over 23 min at a flow of 400 µL/min. The 40 fractions were concatenated into 20 fractions, dried, and reconstituted in 3% acetonitrile and 0.1% trifluoroacetic acid.

### 2.13 nanoLC-MS/MS analysis and database search

Each fraction was analyzed on an Orbitrap Lumos Tribrid mass spectrometer equipped with a FAIMS Pro ion mobility system interfaced with an nLC 1200 liquid chromatography system (all from Thermo Fisher Scientific). Peptides were trapped on an Acclaim Pepmap 100 C18 trap column (100 μm × 2 cm, particle size 5 μm, Thermo Fischer Scientific) and separated on an in-house-constructed analytical column (350 mm × 0.075 mm I.D.) packed with 3 μm Reprosil-Pur C18-AQ particles (Dr. Maisch, Ammerbuch-Entringen, Germany) using a gradient from 3% to 80% acetonitrile in 0.2% formic acid over 85 min at a flow of 300 nL/min. FAIMS Pro alternated between the compensation voltages of –40 and –60, and essentially the same data-dependent settings were used at both compensation voltages. Precursor ion mass spectra were acquired at 120,000 resolution, scan range 450–1375 m/z, and maximum injection time 50 ms. MS2 analysis was performed in a data-dependent mode, where the most intense doubly or multiply charged precursors were isolated in the quadrupole with a 0.7 m/z isolation window and dynamic exclusion within 10 ppm for 60 s. The isolated precursors were fragmented by collision-dissociation at 35% collision energy with a maximum injection time of 50 ms for 3 s (‘top speed’ setting) and were detected in the ion trap, followed by multinotch (simultaneous) isolation of the top-10 MS2 fragment ions within the m/z range 400–1400, fragmentation (MS3) by higher-energy collision dissociation at 65% collision energy and detection in the Orbitrap at 50,000 resolution m/z range 100–500 and a maximum injection time of 105 ms.

The data files for each set were merged for identification and relative quantification using Proteome Discoverer version 2.4 (Thermo Fisher Scientific). The search was against Homo Sapiens (Swissprot Database Mars 2022) using Mascot 2.5 (Matrix Science) as a search engine with a precursor mass tolerance of 5 ppm and a fragment mass tolerance of 0.6 Da. Tryptic peptides were accepted with zero missed cleavage, variable modifications of methionine oxidation and fixed cysteine alkylation, and TMT-label modifications of the N-terminus and lysine. Percolator was used for PSM validation with the strict FDR threshold of 1%. TMT reporter ions were identified with a 3 mmu mass tolerance in the MS3 higher-energy collision dissociation spectra, and the TMT reporter abundance values for each sample were normalized on the total peptide amount. Only the quantitative results for the unique peptide sequences with a minimum SPS match of 50% and an average of signal-to-noise ratio (S/N) above 10 were taken into account for the protein quantification. The reference samples were used as the denominator for calculating the ratios. The quantified proteins were filtered at 1% FDR and grouped by sharing the same sequences in order to minimize redundancy.

### 2.14 Bioinformatics and statistical analysis

Where appropriate, the data in the graphs are reported as the mean ± standard deviation. The statistical analysis was carried out using the GraphPad Prism 6 software (GraphPad Software Inc., La Jolla, CA), and non-paired Student’s t-test was used for multiple comparisons.

To analyze the proteomic data, we used the Qlucore Omics Explorer software (Qlucore, Lund, Sweden) for the principal component analysis, the multi-group comparison, and the unsupervised hierarchical clustering. We used the open access analysis software Funrich ^35^ to compare the proteins listed in the Vesiclepedia ^36^ and ExoCarta ^37^ databases with our data.

### 2.15 Data availability

We have submitted all relevant data of our experiments to the EV-TRACK knowledgebase (EV-TRACK ID: EV240018) ^38^.

## 3. Results

### 3.1 Cellular composition of human melanoma metastatic tissue

We first determined the cellular composition of the metastatic tissues, which were harvested from either melanoma cutaneous metastases or lymph node metastases. In all the tumoral tissues, three distinct cell populations were identified by FACS through size and granulation gating principles. Live cells (aquaDL-negative) were primarily gated based on the expression of CD45 to discriminate leucocytes from non-immune cells (CD45^−^). To better understand the nature of the different identified live cells, we gated on the expression of CD3 (T cells) in the leucocyte group, while the cells belonging to the CD45^−^ gate were identified with a monoclonal antibody (PE) against SOX10 (a melanoma marker) ^39^ (Fig. 2A). The results showed that most of the cells had high side scatter (SSC) and forward scatter (FSC) values (40–80%) indicating that the cells were large and with high intracellular complexity. Moreover, they were negative for CD45 and CD3 but positive for SOX10 (Fig. 2B), suggesting that this subpopulation of cells are tumor cells. The other two subpopulations of cells that were smaller and less granulated were present in similar percentages in the two types of tissues (10–30%). One of these cell populations was identified as T-lymphocytes because these cells were CD45^+^ and CD3^+^, and this population accounted for approximately 15% of the total cells in the lymph node metastases and almost 30% in the skin metastases (Fig. 2C). A mixture of different non-lymphatic cell types was part of the third subpopulation, which generally expressed substantially fewer of the three assessed markers (Fig. 2D). The CD45^−^ and SOX10^−^ populations likely represented other cells of the tumor microenvironment, such as tumor-associated fibroblasts. Overall, the observed representation of the different cell populations was in line with our previous study ^14^ and was confirmed by Hemacolor Rapid staining of cytospin samples. As expected, in the skin metastatic samples the tumor cells as well as macrophages and neutrophils were visible (Fig. 2E). On the other hand, in the lymph node metastatic tissues a large number of cells with lymphocytic morphology was identified, along with macrophages and neutrophils, in parallel with tumor cells (Fig. 2F).

**FIGURE 2.**
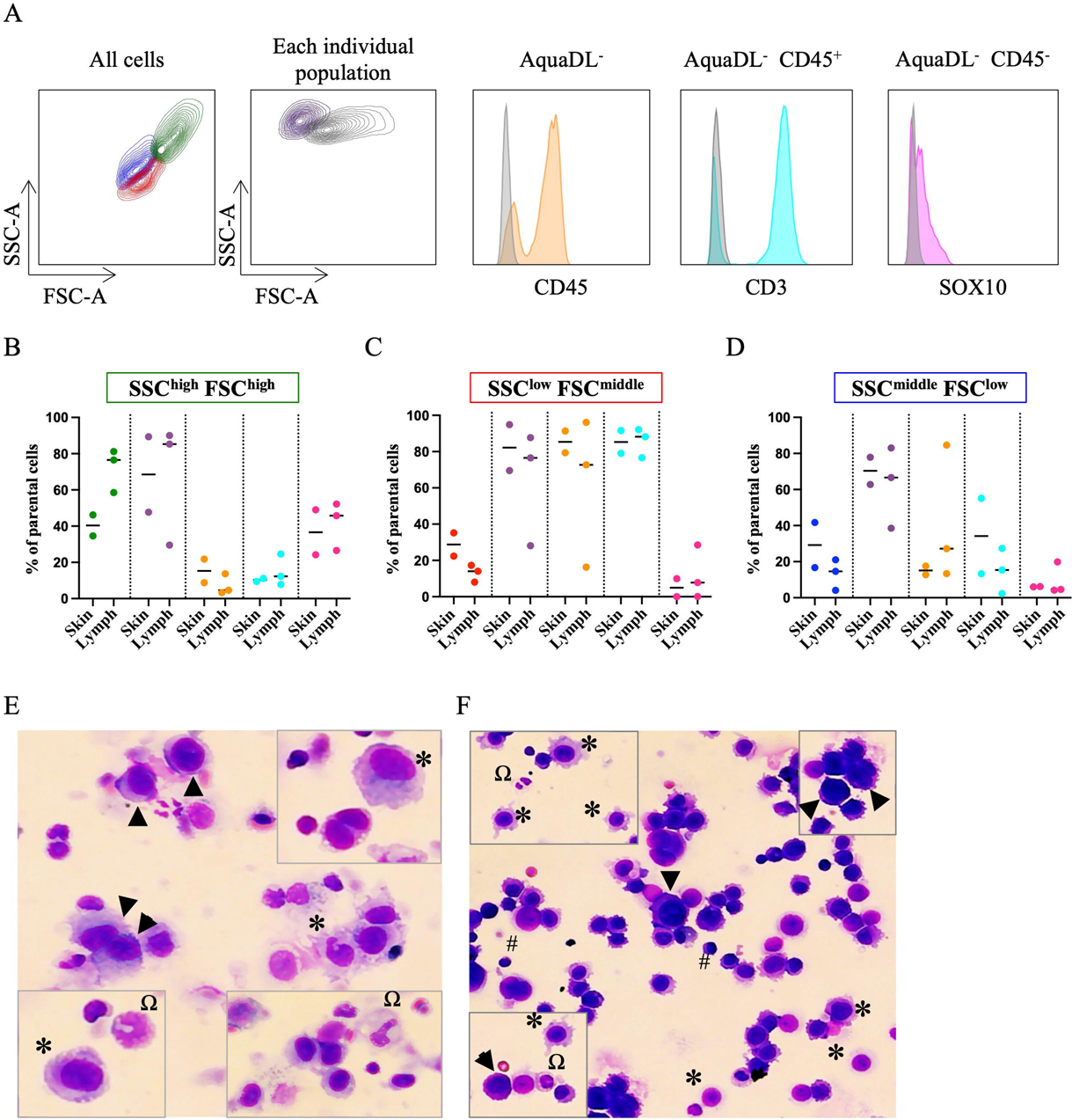
Cellular composition of tumoral samples. (A) Flow cytometry analysis showed that three distinct cell populations were identifiable in tumor tissue samples according to the SSC-A log and FSC-A log, including an SCC^low^/FSC^middle^ subpopulation (red, B); a population rich in large and granulated cells, with high values both for SSC and FSC (green, C); and an SSC^middle^/FSC^low^ population (blue, D). In the different columns of the graphs (B-D) are shown (from left to right) the percentage of the specific cell population out of the total number of parent cells (red, green or blue according to the subpopulation), the percentage of viable cells (purple), and the percentage of cells positive for CD45 (orange), for CD3 (turquoise), and for SOX10 (violet), N = 5. Representative pictures of Hemacolor Rapid staining of the single cell-suspension of (E) melanoma skin metastasis and (F) lymph node metastasis. Black arrows indicate tumor cells; * indicates macrophages; Ω indicates neutrophils; and # indicates lymphocytes. N = 5.

### 3.2 Characterization of EVs isolated from fresh and dry ice-frozen melanoma tissue

After the evaluation of the tissue cellular composition, we isolated and characterized EVs from both fresh and frozen melanoma metastatic tissues as described in Fig. 1. The TEM images showed that isolated EVs were similar from both fresh and frozen tissues and exhibited typical EV morphology and size (30–800 nm) in both conditions (Fig. 3A, Supplementary Fig. 1). Typical EV markers and contaminants were investigated in EV samples isolated from both fresh and frozen tissues as suggested by the MISEV guidelines ^40, 41^. Typical EV markers like CD9, CD63, CD81, and Flotillin-1 were detected in EVs isolated from both fresh and frozen tissues, while Mitofilin was only positive in EVs isolated from patient 3 (Fig. 3B). In addition, the presence of ADAM10, previously reported to be enriched in melanoma-derived small EVs compared to large EVs ^14^, was detected in all of the analyzed samples. Calnexin, an endoplasmic reticulum marker, was positive in both fresh and frozen samples. Even if EVs isolated from two of the patients did not express the investigated markers at the same level, the signal of each marker showed a comparable trend in terms of antigen presence and strength between the EVs from fresh and frozen tissue collected from the same patient. These results indicated a lower intra-group variability compared to inter-patient variability, thus suggesting that EVs isolated from fresh and frozen tissues have similar compositions.

**FIGURE 3.**
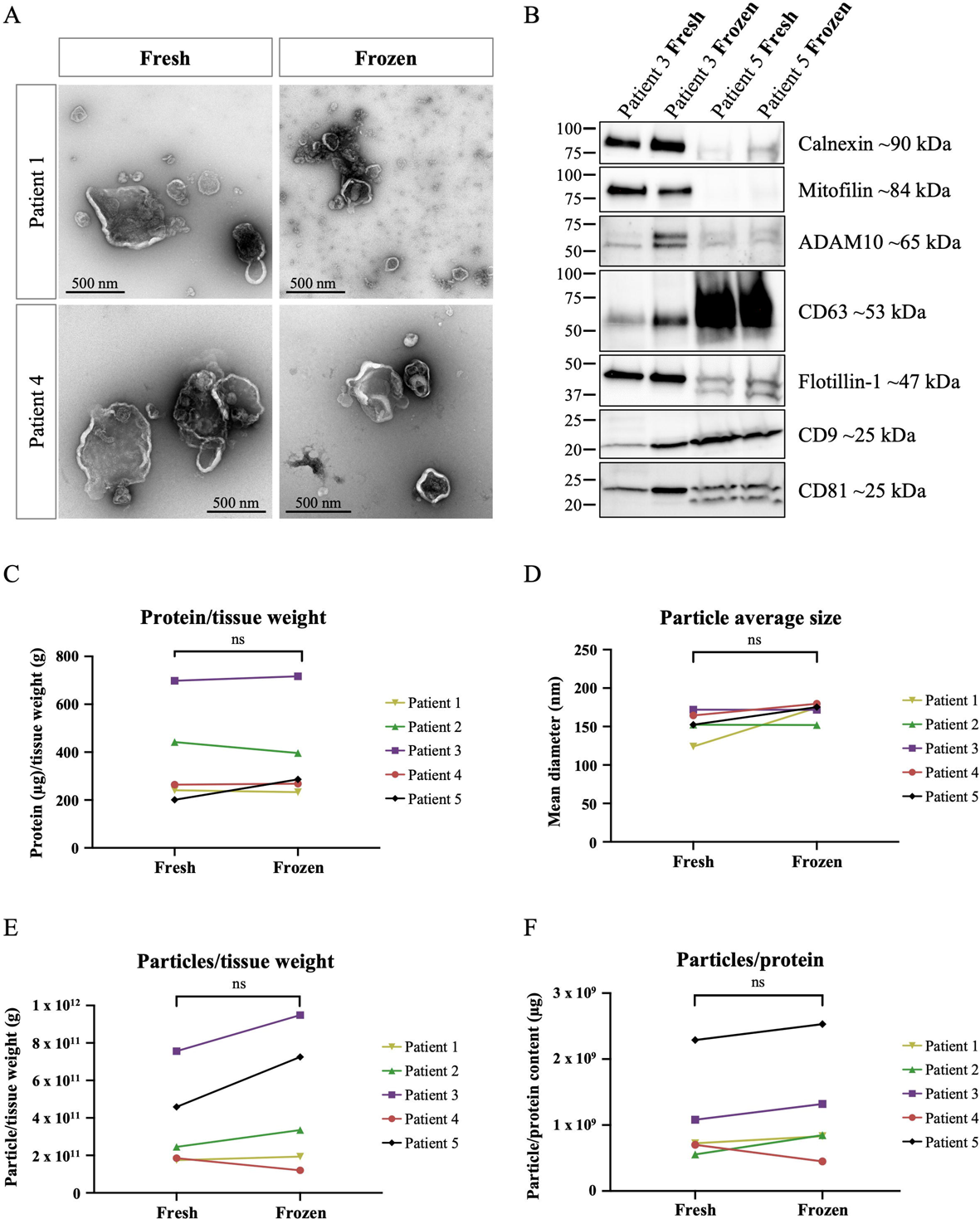
Characterization of the EVs isolated from the fresh and frozen melanoma tissues. (A) Representative microphotographs of EVs isolated from fresh and frozen melanoma tissues. Scale bar: 500 nm. N = 2. (B) Western blot performed on both experimental groups from two patients to evaluate the presence of EV markers such as CD9, CD63, CD81, and Flotillin-1 as well as Mitofilin, ADAM10, and Calnexin, an ER-derived protein. N = 2. The similarity between EVs from the fresh and frozen tissue samples was assessed and compared in terms of (C) protein/tissue weight ratio, (D) particle average size, (E) particles/tissue weight ratio, and (F) particles/protein ratio. ns = non-significant. N = 5.

A crucial point to determine whether frozen tissues are a valid source of EVs is to verify if and to what extent the freezing process affects the yield and the purity of the EV preparation. We first quantified the protein concentration in the EVs isolated from the two experimental groups, and we normalized the data to the weight of the tissue collected from each patient. The ratio between the EV protein concentration and tissue weight was comparable in the EVs isolated from the fresh and the frozen tissues (Fig. 3C). Similarly, the average particle diameter was not significantly different between fresh and frozen samples within the same patient (Fig. 3D). We further analyzed the ratio between the number of particles and the weight of the tissue. As shown in Fig. 3E, the number of particles per tissue weight was slightly higher in EVs isolated from the frozen samples compared to those obtained from the fresh ones, indicating that some non-EV contamination may appear in the frozen samples. Only the samples from skin metastasis showed an opposite trend, and in patient 1 the particles to weight ratio was substantially the same for fresh and frozen tissues, while in patient 4 the ratio was lower in the frozen group. Importantly, despite this observed variability, no statistically significant differences were observed comparing fresh and frozen samples. Lastly, to quantify the purity of EV preparations we calculated the ratio between the particle number and the protein concentration. Although in patient 4 the ratio was slightly lower in frozen tissue than in fresh tissue, overall no significant differences were evident between the two experimental groups using this approach (Fig. 3F).

### 3.4 Quantitative proteomics analysis of EVs isolated from fresh and frozen tissue

Next, we performed quantitative TMT proteomics analysis of EVs isolated from fresh and frozen tissue to further investigate the EV composition and to evaluate the similarity within the two groups, as well as to evaluate signs of potential cell-derived contamination. A total of 8251 proteins were identified, and of these 7573 were quantified (all proteins are listed in Supplementary Table 1).

First, we performed a principal component analysis (PCA) on the quantified proteins to visualize the relation between the different groups of EV samples. The PCA plot showed that the samples were more similar for EVs isolated from the same patient than for the EVs isolated from the tissues treated in the same condition (fresh or frozen) (Fig. 4A). The same PCA plot is shown in Suppl. Fig 2A but with coloring based on fresh vs. frozen tissue-derived EVs instead of coloring based on patients. These results indicate that the intra-group variability of the protein cargo was lower compared to the inter-patient variability, again suggesting that the tissue freezing did not greatly alter the protein composition of the isolated EVs.

**FIGURE 4.**
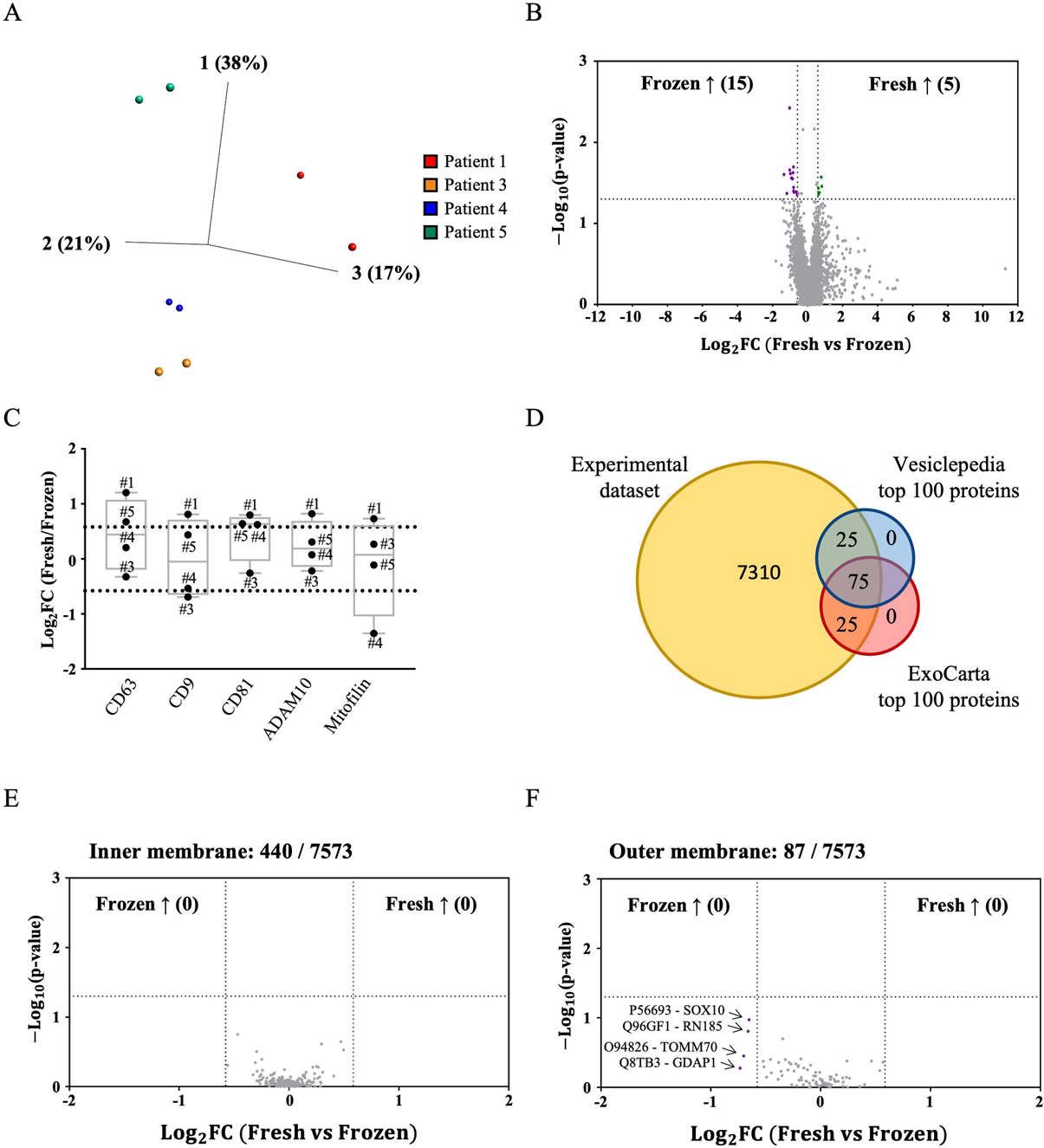
Quantitative proteomic analysis (TMT) performed on the fresh and frozen tissue-derived EVs from four patients. (A) PCA illustrating the relation between EVs isolated from different patients. (B) The volcano plot shows the 15 and 5 proteins that were significantly enriched in frozen (purple) and in fresh (green) tissue-derived EVs, respectively. N = 4. (C) Fold change values of EV markers (CD63, CD9, CD81, ADAM10, Mitofilin) in fresh versus frozen samples. N = 4. (D) Venn diagram showing the comparison of the whole protein dataset obtained from the four patients (yellow) and the Vesiclepedia (blue) ^36^ and ExoCarta (red) ^37^ databases. Volcano plots of the inner (E) and outer (F) membrane mitochondrial proteins found in the experimental dataset.

The small differences in protein content in fresh and frozen tissue-derived EVs were confirmed by volcano plot analysis (Fig. 4B). Only 15 proteins were significantly enriched in the frozen samples (listed in Supplementary Table 2), while 5 were significantly enriched in the fresh samples (listed in Supplementary Table 2). In addition, only 8 proteins showed a greater than 4-fold change in the fresh sample compared to the frozen sample (Fig. 4B), although none of these differences were significant (Supplementary Table 3). Moreover, the proteomics results did not show any significant enrichment of intracellular contaminants in the frozen samples.

To further determine if the tissue freezing procedure affected our isolated EVs, we compared the relative abundance of five proteins (CD63, CD9, CD81, ADAM10, and Mitofilin) previously found to be enriched in EVs ^14, 42^. The fold changes for the five proteins were between 1.2 and 1.3 indicating no major difference in the abundance of these proteins in isolates from fresh or frozen tissues (Fig. 4C). In addition, our dataset contained all the top 100 most-represented proteins in the Vesiclepedia ^36^ (http://microvesicles.org/) and ExoCarta ^37^ (http://exocarta.org/index.html) databases. In particular, 75 of these most represented proteins were in common between our datasets and both the Vesiclepedia ^36^ and ExoCarta ^37^ lists, while 25 were shared between our dataset and one of the online databases (Fig. 4D). This comparison showed that most of the proteins found in our samples are associated with EVs because they have been found in EVs in previous studies. Altogether, these results demonstrated that tissue-derived EVs were similar in protein cargo when isolated from both fresh and frozen tissues.

We then further investigated any presence of potential intra-cellular contaminations in tissue-derived EV isolates, especially focusing on mitochondrial proteins because these can be a sign of cell disruption after freezing. We identified 440 proteins associated with the mitochondrial inner membrane, and none of them showed a significative enrichment in the frozen samples compared to the fresh samples (Fig. 4E). On the other hand, 87 outer membranes proteins were detected, and among these four showed a small and non-significant enrichment in the frozen samples (Fig. 4F and Supplementary Table 4). Of note, one of these proteins was the transcription factor SOX-10, which has been proposed to be a marker of melanoma tumors ^39, 43^ and is what we used to detect tumoral cells in the flow cytometry analysis.

### 3.5 Evaluation of the potential diagnostic value of fresh and frozen tissue-derived EVs

Lastly, we wanted to determine whether the EVs isolated from frozen tissue keep their potential diagnostic value. To do that, we compared the relative abundance from the mass spectrometry analysis of five proteins (MT-CO2, COX6c, SLC25A22, HLA-DR, and Erlin2) that we have previously shown to be enriched in EVs isolated from fresh human melanoma tissue and in the plasma of patients affected by melanoma, breast cancer, and ovarian cancer ^12^. We did not find any significant differences in the levels of the five analyzed markers between fresh and frozen tissue-derived EVs. However, slightly greater differences were observed in EVs isolated from skin metastases (patient 1 and 4) compared to lymph node metastases (patient 3 and 5) (Fig. 5A). ELISA assays and Western blot confirmed the presence of two of these proteins in both fresh and frozen tissues (Fig. 5B-E). These findings showed that the EVs isolated from frozen tissues retain a similar potential diagnostic value to those obtained from fresh specimens.

**FIGURE 5.**
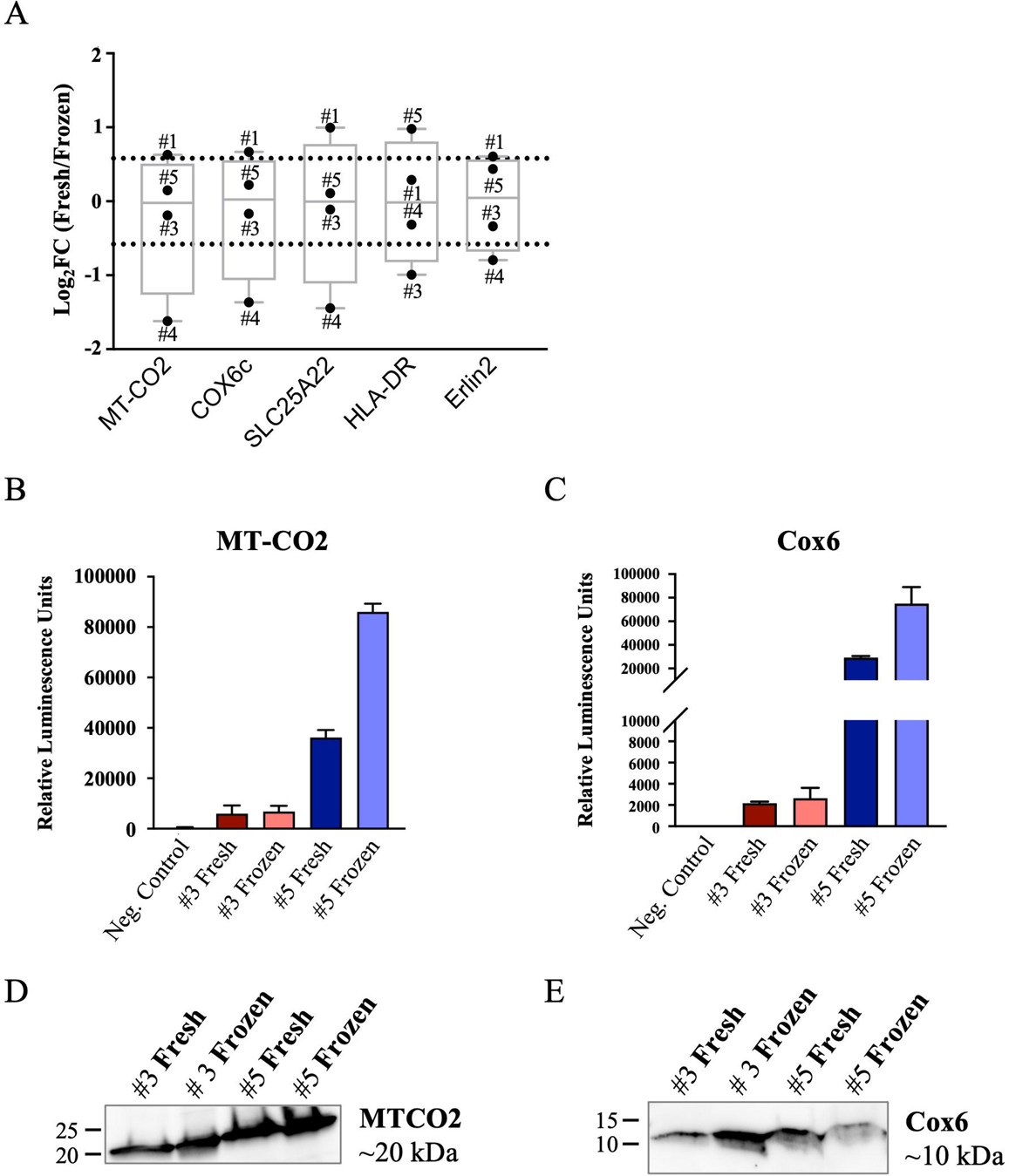
Evaluation of the potential diagnostic value of fresh and frozen tissue-derived EVs. (A) The relative quantity of five potential EV-melanoma markers (MT-CO2, COX6c, SLC25A22, HLA-DR and Erlin2) was determined from the proteomic analysis performed on the two experimental groups. The data are reported as the average fold change in the ratio between the fresh and frozen tissue-derived EVs. N = 4. ELISA assay of (B) MT-CO2 and (C) Cox6 levels in EVs isolated from fresh and frozen tissues from patients 3 and 5. EVs from HEK293F cells were included as the negative control. N = 2. Western blot analysis of (D) MT-CO2 and (E) Cox6 levels in EVs derived from fresh and frozen tissues from patients 3 and 5. N = 2.

## 4. Discussion

Although in previous studies both fresh ^12, 14, 15, 44–46^ and frozen ^13, 16, 17, 25–33^ tissues have been used as sources of tissue-derived EVs, there is still the unmet need to understand if and to what extent the freezing might affect the purity or composition of isolated EVs. In this study we demonstrate that the EVs isolated from frozen tissues are similar to the EVs isolated from fresh tissues in terms of morphology, yield, and protein composition. Importantly, we show that EVs isolated from frozen tissues maintain a protein composition that we previously showed in fresh tissue to include potential tumor biomarkers ^12^. This finding highlight that the freezing process does not seem to significantly contaminate tissue-EV isolates with non-vesicular material.

First, we characterized the cellular composition of melanoma tissues through flow cytometry analysis. Our results were consistent with a previous report, with similar relative numbers of white blood cells and presumed melanoma cells ^14^. However, we observed SOX10^+^ cells in all melanoma metastatic tissues analyzed, indicating the presence of tumor cells in the tissues ^39, 43^. Moreover, quantitative proteomic analysis showed the presence of SOX10 in EVs from both fresh and frozen tissues with a small and non-significant enrichment in the frozen samples. The presence of SOX10 in melanoma-derived EVs could make the SOX10^+^ EVs a potential marker of melanoma. However, further validation studies are needed to determine the potential role of this vesicular protein as a true melanoma biomarker.

The comparison of TEM micrographs of EVs isolated from fresh and frozen melanoma tissues showed no visible differences or the presence of debris between the fresh and frozen tissues. The similarity between EVs isolated from fresh and frozen tissues was confirmed by quantitative proteomic analysis showing no significant protein enrichment in any type of sample. This is in line with a previous study comparing EVs from epithelial ovarian cancer that were harvested either from fresh or frozen tissue ^47^. Of note, all of the top-100 proteins annotated in the two main EV databases (Vesiclepedia ^36^ and Exocarta ^37^) were found in EVs from both fresh and frozen tissues, further supporting the EV origin of samples from both types of tissue (Fig. 4D). Only 15 proteins were significantly enriched in the frozen group (Suppl. Table 2), and all of them were listed in Vesiclepedia. However, they belong to intracellular compartments, suggesting that they may potentially be contaminants in the EV isolates from frozen tissue.

Even though we used exclusively human melanoma metastatic tissues as the source of EVs, we processed two different types of metastases, from either skin or lymph nodes. The two sample types have very different textures and compositions prior to EV isolation. However, we did not observe any significant difference in morphology, concentration, or protein composition in EVs isolated from skin melanoma metastases versus lymph node metastases. Our results support the idea that the dry ice freezing procedure is a reasonable way to preserve EVs in tissues for future isolation, regardless of the melanoma metastasis tissue origin. However, we exclusively used human melanoma tissue, which should be considered by any researcher aiming at isolating EVs from other types of frozen tissues.

We isolated large and small EVs by serial ultracentrifugation and combined both EV subpopulations before performing an iodixanol density cushion centrifugation to remove potential contaminants. Because we did not keep the EV subpopulations separated, we cannot determine if any of the proteins found in the EV samples were actually enriched in one the EV subpopulations. Moreover, we cannot exclude that one or more specific EV subpopulations might be partially lost during the freezing process. Which EV subpopulation is preferable for use as a disease biomarker is still a matter of discussion, so it would be interesting to explore if the freezing process has any effects on specific EV subpopulation. Moreover, only one way to freeze the samples was evaluated, and we chose to test the most commonly used method in biobanks. It would be interesting to evaluate the characteristics of EVs isolated from samples frozen in different ways because it has been demonstrated that different freezing approaches have different effects on tissues ^20^ and might contribute with different types and degrees of contaminants and thus have disparate effects on the EV composition. Moreover, we saved the frozen samples at –80°C for a relatively short time (two weeks) compared to the storage time of samples in the biobank (months or years). Further experiments are needed to analyze EVs isolated from samples frozen for longer times with the aim of definitively demonstrating that tissues saved in a biobank can be used as a valid source of EVs.

The relative absence of cell-derived contamination in the EVs isolated from the frozen tumor tissues, and the consequent similarity with the fresh counterpart, was further supported by the analysis of mitochondrial proteins, which can identify cell damage induced by things like freezing. However, mitochondrial proteins are known to be physiologically and actively packaged within EVs ^48, 49^, and we previously reported that proteins of the inner mitochondrial membrane can potentially serve as EV-associated biomarkers of melanoma ^12^. In addition, in some pathological cases whole mitochondria have been reported to be enclosed within large EVs that originate from the endosomal pathway ^50^. However, we did not identify any significant increase of mitochondrial proteins in the frozen tissue derived-EVs. Interestingly, among the proteins of the outer mitochondrial membrane the already mentioned transcription factor SOX10 was detected in both frozen and fresh tissue-derived EVs. Because this marker is considered to be highly specific for melanoma tumors ^39, 43^, we suggest that it could potentially represent an EV melanoma marker.

The data we present support the use of frozen tissue-derived EVs for diagnostic purposes because the levels of all the potential biomarkers that were assessed were similar to those found in the fresh tissue-derived EVs. This was not surprising considering that EVs can protect the biomolecules they carry from the potential damage of the freezing step ^51^. Indeed, it was previously reported that the content in terms of mRNA and miRNA, molecules that are much less stable than proteins, in EVs isolated from frozen tissues was comparable to that of the fresh tissue-derived EVs ^47^. These results were in contrast with those obtained from cells, in which gene expression changed drastically in frozen postmortem brain samples compared to fresh samples ^52^.

Overall, the results described in this work suggest that the frozen tissues already available in biobanks might be used as sources for EV-based biomarker research. This would expand and facilitate the study in the field of EVs from solid tissues and their potential diagnostic role by increasing the amount of starting material available for research.

## 5. Conclusions

Through a deep quantitative proteomic analysis of EVs isolated from both fresh and frozen melanoma tissues, we demonstrate that the freezing process using dry ice does not significantly affect the protein composition of EVs isolated from tissues. Consequently, we have demonstrated the effectiveness of the described protocol for isolating EVs from frozen tissue, as previously optimized by our group using fresh tissues ^12, 14, 15^. Moreover, the results support the potential use of the EVs from frozen tissues as a source of disease-associated biomarker discovery.

## Acknowledgment

We acknowledge the Centre for Cellular Imaging at University of Gothenburg and the National Microscopy Infrastructure (VR-RFI 2019-00217) for providing assistance in microscopy. For proteomic analysis we thank Annika Thorsell at the Proteomics Core Facility at Sahlgrenska Academy, University of Gothenburg. The proteomic analysis was performed with financial support from SciLifeLab and BioMS.

## FIGURE LEGENDS

**SUPPLEMENTARY FIGURE 1** Representative close-up electron micrographs of EVs isolated from two patients from both fresh and frozen tissues. Scale bar: 100 nm. N = 2.

**SUPPLEMENTARY FIGURE 2** PCA illustrating the relation between fresh (red) and frozen (violet) tissue-derived EVs. The samples belonging to the same patient are surrounded by a circle.

**SUPPLEMENTARY TABLE 1** List of proteins quantified in EVs isolated from fresh and frozen tissue. N = 4

**SUPPLEMENTARY TABLE 2** Significant proteins identified and quantified in EVs from fresh versus frozen tissues. The list shows the significant (p-value < 0.05) proteins that were enriched or reduced with a fold change greater than 1.5.

**SUPPLEMENTARY TABLE 3** The list shows the proteins that were enriched in EVs from fresh samples with a fold change > 16 but not significantly different (p-value > 0.05).

**SUPPLEMENTARY TABLE 4** The list shows the outer mitochondrial proteins that were enriched in EVs from frozen samples with a fold change > 1.5 but not significantly different (p-value > 0.05).

