## Supplementary figures and images for "Extracellular vesicles isolated from frozen and fresh human melanoma tissue are similar in purity and protein composition"

### Supplementary Fig 1

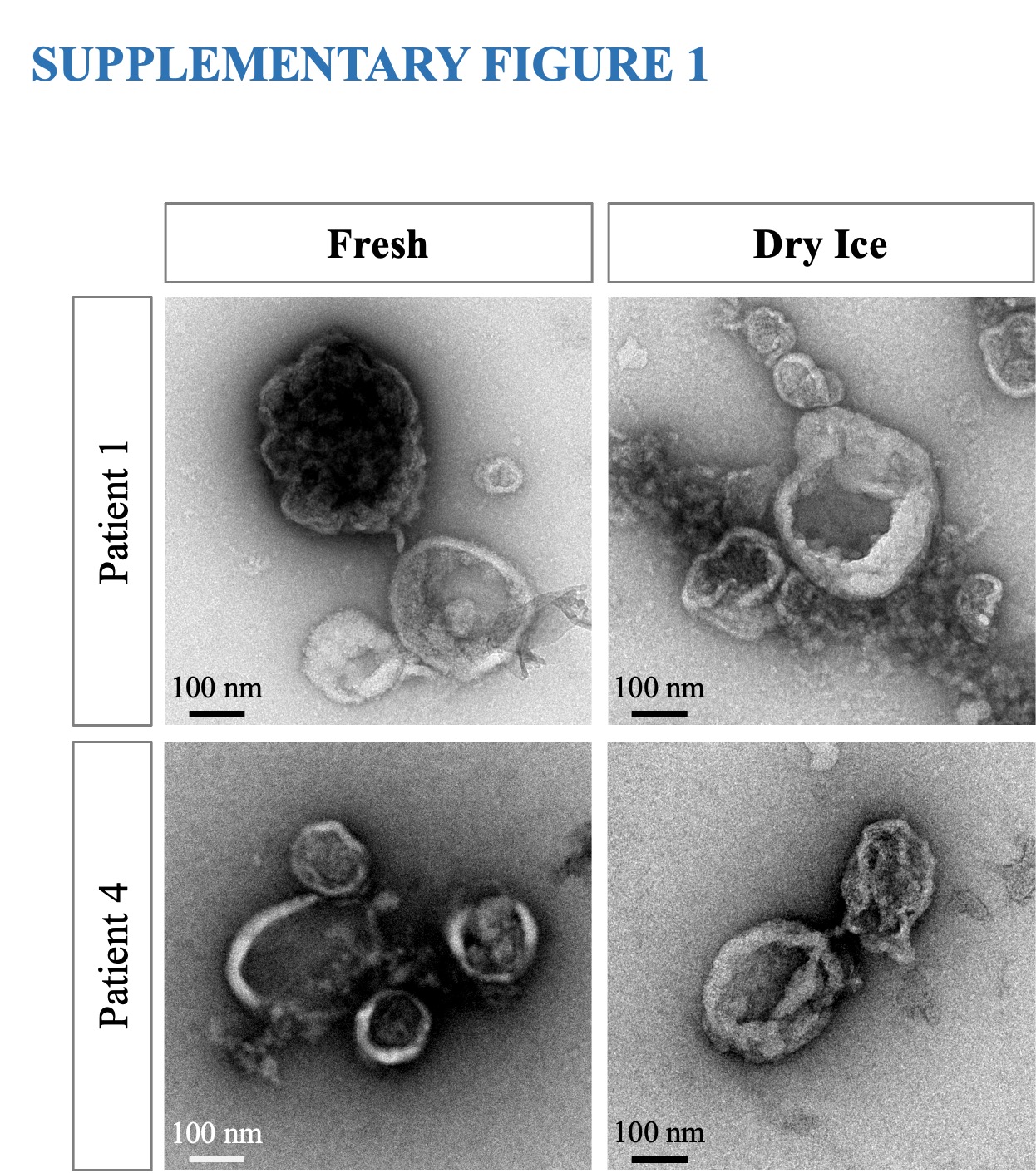

### Supplementary Fig 2

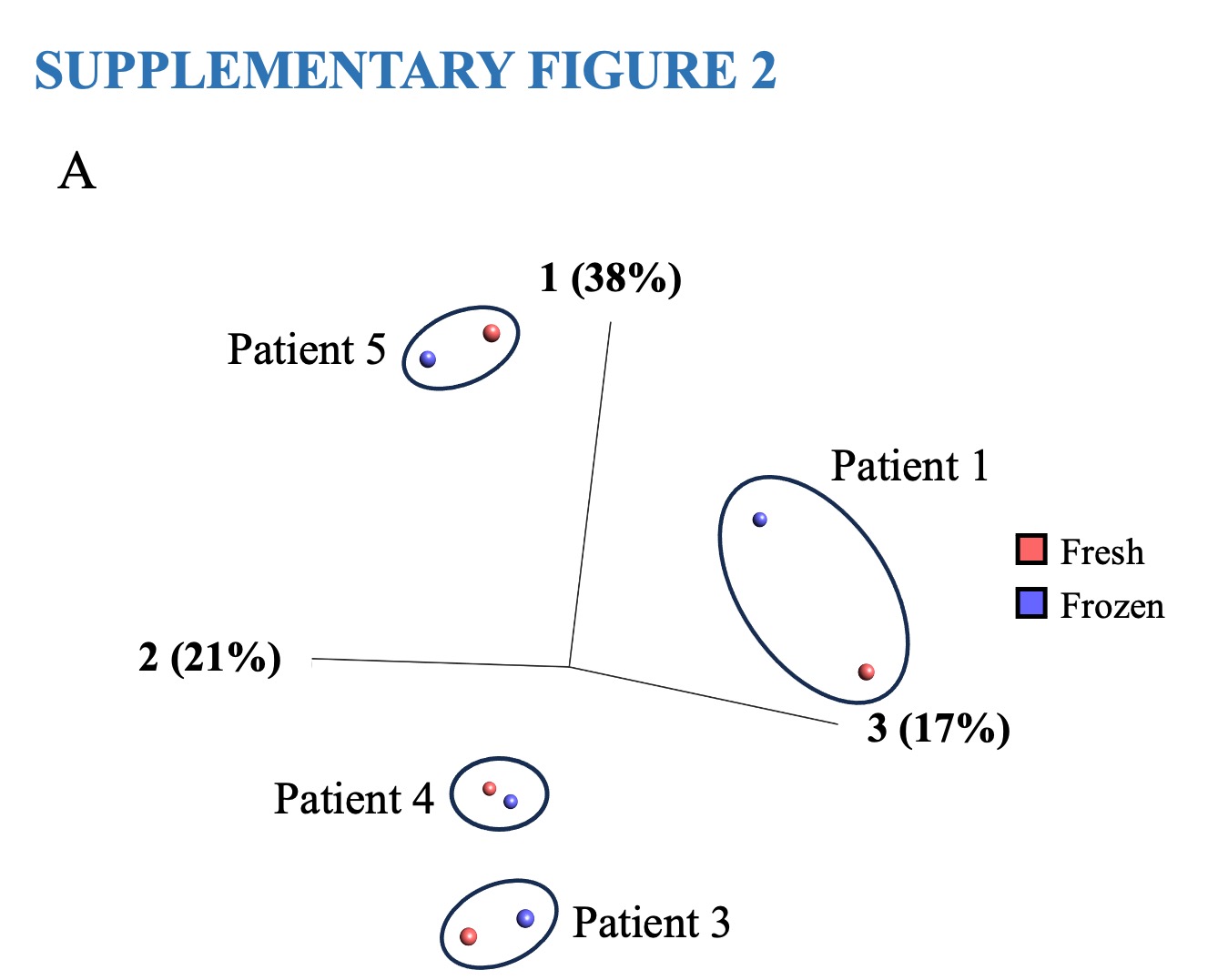
