## Supplementary Table 2 for "Extracellular vesicles isolated from frozen and fresh human melanoma tissue are similar in purity and protein composition"

**Table S2:** Significant proteins identified and quantified in EVs from fresh versus frozen tissues.

The list shows the significant proteins (p-value < 0.05) that were enriched or reduced with a fold change larger than 1.5. Fold change means fresh tissue-EVs / frozen tissue-EVs.

| **Accession ID** | **Description** | **Gene name** | **Log_2_ (Fold change)** |
| --- | --- | --- | --- |
| P18615 | Negative elongation factor E | NELFE | **-1.344** |
| Q8TF01 | Arginine/serine-rich protein PNISR | PNISR | **-1.185** |
| Q15527 | Surfeit locus protein 2 | SURF2 | **-1.024** |
| Q9UHA4 | Ragulator complex protein LAMTOR3 | LAMTOR3 | **-1.022** |
| Q6PK04 | Coiled-coil domain-containing protein 137 | CCDC137 | **-0.979** |
| Q14BN4 | Sarcolemmal membrane-associated protein | SLMAP | **-0.924** |
| Q01105 | Protein SET | SET | **-0.864** |
| P0DME0 | Protein SETSIP | SETSIP | **-0.864** |
| Q9BQE4 | Selenoprotein S | SELENOS | **-0.828** |
| Q9UBT2 | SUMO-activating enzyme subunit 2 | UBA2 | **-0.806** |
| Q01081 | Splicing factor U2AF 35 kDa subunit | U2AF1 | **-0.800** |
| O95232 | Luc7-like protein 3 | LUC7L3 | **-0.799** |
| Q8NFW8 | N-acylneuraminate cytidylyltransferase | CMAS | **-0.772** |
| Q69YL0 | Protein NCBP2AS2 | NCBP2AS2 | **-0.651** |
| P67809 | Y-box-binding protein 1 | YBX1 | **-0.596** |
| Q14168 | MAGUK p55 subfamily member 2 | MPP2 | **0.810** |
| Q96K17 | Transcription factor BTF3 homolog 4 | BTF3L4 | **0.777** |
| Q9Y6E0 | Serine/threonine-protein kinase 24 | STK24 | **0.674** |
| Q9Y5Y0 | Feline leukemia virus subgroup C receptor-related protein 1 | FLVCR1 | **0.625** |
| P15289 | Arylsulfatase A | ARSA | **0.619** |
