## Supplementary Table 3 for "Extracellular vesicles isolated from frozen and fresh human melanoma tissue are similar in purity and protein composition"

**Table S3:** List of the 8 proteins enriched in EVs from fresh versus frozen tissues.

The list shows the proteins that were enriched in the fresh-derived EV samples with a mean fold change > 16 but not significantly different (p-value > 0.05).

| **Accession ID** | **Description** | **Gene name** | **Subcellular localization** | **FC_#1** | **FC_#3** | **FC_#4** | **FC_#5** | **FC_Mean** | **P-value** |
| --- | --- | --- | --- | --- | --- | --- | --- | --- | --- |
| Q15323 | Keratin | KRT31 | NA | 0.017 | 27.8 | 10000 | 0.239 | 2507.014 | 0.362 |
| P17028 | Zinc finger protein 24 | ZNF24 | Nucleus | 1.093 | 1.712 | 113.6 | 0.747 | 29.288 | 0.641 |
| Q96AZ6 | Interferon-stimulated gene 20 kDa protein | ISG20 | Nucleus | 0.362 | 0.886 | 67.3 | 0.726 | 17.319 | 0.527 |
| Q5TF58 | Intermediate filament family orphan 2 | IFFO2 | NA | 0.031 | 0.635 | 93.2 | 1.166 | 23.758 | 0.632 |
| Q9Y312 | Protein AAR2 homolog | AAR2 | NA | 0.887 | 0.332 | 127.1 | 0.343 | 32.166 | 0.632 |
| Q0D2I5 | Intermediate filament family orphan 1 | IFFO1 | NA | 0.031 | 0.635 | 93.2 | 1.166 | 23.758 | 0.632 |
| P01714 | Immunoglobulin lambda variable 3-19 | IGLV3-19 | Secreted | 1.091 | 0.527 | 72.3 | 1.222 | 18.785 | 0.879 |
| Q6ZMN7 | PDZ domain-containing RING finger protein 4 | PDZRN4 | NA | 1.133 | 0.859 | 136.3 | 0.728 | 34.755 | 0.5 |
