## Supplementary Table 4 for "Extracellular vesicles isolated from frozen and fresh human melanoma tissue are similar in purity and protein composition"

**Table S4:** List of the 4 mitochondrial proteins enriched in EVs from frozen versus fresh tissues.

The list shows the mitochondrial outer membrane proteins that were enriched in EVs from frozen tissues with a fold change > 1.5 but not significantly different (p-value > 0.05).

| **Accession ID** | **Description** | **Gene name** | **Log_2_ (Fold change)** | **P-value** |
| --- | --- | --- | --- | --- |
| O94826 | Mitochondrial import receptor subunit TOM70 | TOMM70 | -0.701 | 0.353 |
| Q8TB36 | Ganglioside-induced differentiation-associated protein 1 | GDAP1 | -0.734 | 0.53 |
| Q96GF1 | E3 ubiquitin-protein ligase RNF185 | RNF185 | -0.660 | 0.156 |
| P56693 | Transcription factor SOX-10 | SOX10 | -0.652 | 0.107 |
